# Infrared thermography for plant stress detection in vertical farms: Investigating spatiotemporal variations and exploring solutions via machine learning

**DOI:** 10.1101/2024.07.04.602094

**Authors:** Avinash Agarwal, Filipe de Jesus Colwell, Rosalind Dinnis, Viviana Andrea Correa Galvis, Tom Hill, Neil Boonham, Ankush Prashar

**Affiliations:** School of Natural and Environmental Sciences, Newcastle University, Newcastle upon Tyne, UK; Crop Science R&D Division Infarm - Indoor Urban Farming B.V., Amsterdam, The Netherlands; Faculty of Medical Sciences, Newcastle University, Newcastle upon Tyne, UK

## Abstract

Application of infrared thermography (IRT) for real-time plant stress detection has grown rapidly in recent years. Although the technology has been well established for crops grown in fields and glasshouses, its feasibility for vertical farms has not been tested extensively. In this study, temporal monitoring of stress induced by root dehydration in purple basil plantlets inside a vertical farm was performed to identify bottlenecks in real-time stress detection via IRT. Subsequently, potential solutions were investigated via machine learning by implementing support vector machines for supervised classification. Edge effects as well as proximity to air vents were identified as the major causes of positional variation in plant temperature that could lead to misprediction of stress. Binary, ternary, and quaternary classification models were trained using thermal images from two, three, and four levels of stress, respectively, to assess model performance. Binary classification models trained with plants experiencing medial and high levels of stress were able to identify stressed plants with high accuracy (81–94%). Further, binary models trained using plants under medial levels of stress generated a continuous probability distribution for stress prediction when plotted against plant temperature. In contrast, models trained using samples experiencing high stress generated distinct probabilistic clusters for the unstressed and highly stressed plants, but were unable to classify medial stress samples reliably. Similarly, ternary and quaternary models were able to better predict very high and very low levels of stress than intermediate stress levels. Hence, our findings suggest that binary classification models trained using samples under medial levels of stress would be helpful in overcoming spatiotemporal variations in canopy thermal profile by providing reliable probabilistic estimates of plant stress within a vertical farming system.

**Key points:**

1. Plant stress detection in vertical farms via thermal imaging may be challenging because perceptible plant temperature can be strongly influenced by its microenvironment.
2. Thermal image analysis via supervised machine learning allows the development of robust prediction models that can overcome such factors to identify stressed plants.
3. Binary classification machine learning models can reliably identify stressed plants as well as provide probabilistic estimates for the degree of stress.

## 1 Introduction

Plant water status is a crucial factor influencing crop growth and yield as it plays an important role in the biosynthesis and transport of nutrients, phytohormones, carbohydrates, and various other metabolites (Cornic and Massacci, 1996; Farooq et al., 2012; Lambers and Oliveira, 2019). Additionally, plant water status also acts as an indicator of stress by influencing temperature regulation via stomatal opening (Mahan and Upchurch, 1988; Jones, 1999). Any biotic or abiotic stress that disrupts water uptake results in the closure of stomata due to loss of turgor pressure, which in turn impedes transpirational cooling of leaves, increasing canopy temperature (Grant et al., 2006; Ben-Gal et al., 2009). This increment in canopy temperature occurs before plant water status is affected markedly, making it a useful trait for early stress detection (Grant et al., 2006; Prashar and Jones, 2014).

Considering this, infrared thermography (IRT) has gained popularity in recent years for real-time monitoring of plant water status and stress (Prashar and Jones, 2014; Waiphara et al., 2022). Since IRT accurately senses fluctuations in canopy temperature, its pertinence in high-throughput real-time crop stress monitoring has been well-explored for crops grown in fields, greenhouses, and growth chambers (Grant et al., 2006; Padhi et al., 2012; Prashar et al., 2013; Prashar and Jones, 2014; Lima et al., 2016; Biju et al., 2018; Khorsandi et al., 2018; Pipatsitee et al., 2018; Pineda et al., 2021). However, rapid increase in plant factories over the past decade has necessitated the translation of this technology to vertical farming systems as well.

Despite better regulation of growth environment within vertical farms (Kalantari et al., 2018), plants are still vulnerable to stress due to biotic or abiotic stressors (Roberts et al., 2020). Further, high-density planting increases the likelihood of rapid widespread crop loss if the stress is not mitigated timely. Since plant growth in vertical farms is extensively automated, with minimal human intervention, introduction of fast high-throughput plant monitoring systems that complement any alerts coming from hardware failures is necessary for ensuring quality production. Thus, IRT may be deemed as an ideal tool for real-time crop monitoring in vertical farms owing to its rapid response, sensitivity, ease of use, and high throughput.

Considering the high dimensionality of thermal images, processing of IRT data may be challenging. However, the process can be simplified by implementing machine learning (ML). In recent years, ML has emerged as a useful tool for high-throughput analysis of image-based datasets as it enables rapid in-depth processing of numerical and spatial data patterns (Singh et al., 2016; Liakos et al., 2018; Gao et al., 2020; Zubler and Yoon, 2020). Amongst the various ML approaches, supervised learning via support vector machines (SVMs) has been frequently employed for detecting plant stress via multispectral, thermal, and hyperspectral imaging (Sankaran et al., 2013; Raza et al., 2014; Calderón et al., 2015; Cao et al., 2018) as it creates hypothetical boundaries in high-dimensional space for optimal separation between different dataset classes (Cortes and Vapnik, 1995). Hence, integrating SVM-based ML classification with IRT could streamline the process for real-time crop monitoring within vertical farms.

In the present study, we have explored the feasibility of IRT for real-time stress detection in vertical farms using purple basil (*Ocimum basilicum* L. var. *purpurascens*) as a model system. Since implementation of IRT for plant stress detection in vertical farms has not been investigated extensively till date, we carried out an exploratory study to reveal potential bottlenecks in this process. For this, we imposed stress on the plants by withholding irrigation to elicit changes in canopy temperature, and monitored the plants at regular intervals via thermal imaging to obtain spatiotemporal trends. The information was used to develop a framework by combining IRT with ML for real-time plant stress detection.

## 2 Methods

### 2.1 Plant material

Purple basil seedlings were raised in coco-peat plugs (Van der Knapp, The Netherlands) in a nursery chamber (Aralab-InFarm UK Ltd., London, UK), each plug containing 4–5 seedlings. At ∼2 cm the seedlings were transferred to an experimental vertical farm (InStore Farm, InFarm UK Ltd.; Figure 1A) at the Agriculture Building, Newcastle University, UK. The farm comprised of eight plant growth trays (80×80 cm^2^), each having an 8×8 array of empty slots (Figure 1B). Each tray received 58 seedling plugs. A commercial hydroponics fertiliser mix was used as the nutrient source, and irrigation was performed following the nutrient film technique using an electrical pump (Figure 1A). A broad-spectrum LED array having approximate red (400–499 nm):green (500–599 nm):blue (600–699 nm) distribution of 40:20:40 was used to provide 280 *µ*mol/m^2^sec PPFD in a 16/8 h day-night cycle. Temperature and relative humidity were maintained at 25±1 °C and 60±5%, respectively, through a centralised air circulation system. Growth conditions and irrigation were monitored on a Farmboard (InFarm® UK Ltd.) using built-in sensors for temperature, humidity, flow rate, electrical conductivity, and pH.

**Figure 1.**
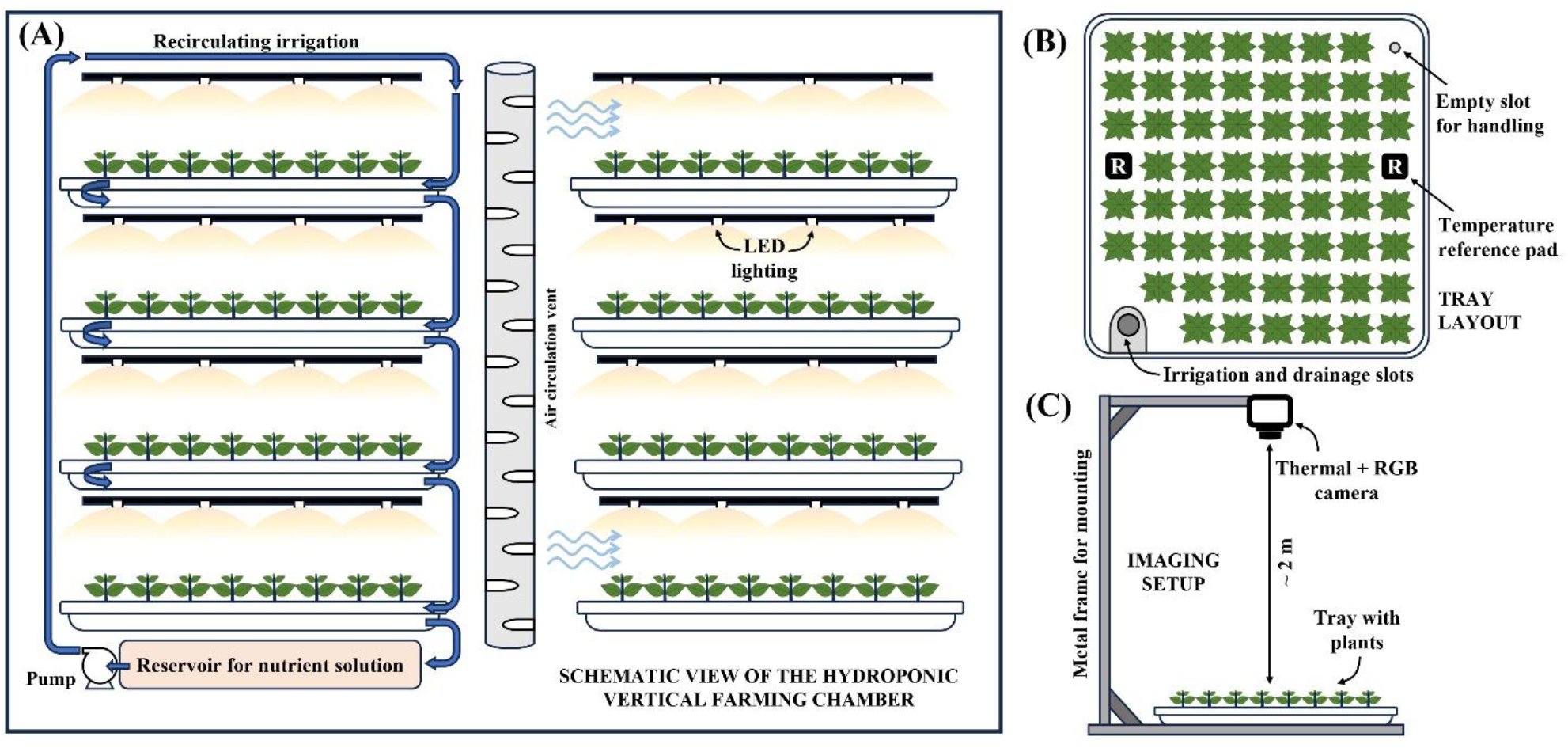
Schematic overview of the experimental vertical farming unit (A), layout of the planting tray (B), and the setup for imaging (C).

### 2.2 Experimental design

Two preliminary trials (PTs) were carried out to get a general overview of stress responses induced by water deficit, followed by a third (main) trial aimed at ML-based stress detection (Table 1) in two-week-old plants. In the first trial (PT-1), irrigation was stopped at 8 AM, and thermal imaging (described later) was carried out with five separate trays for different durations without irrigation as follows: for 0, 6, and 12 h without irrigation on the same day, and for 22 and 34 h without irrigation on the next day. Twelve samples were randomly selected from the corresponding tray immediately after imaging, and the plants were cut at the base of the stems. The aerial part was considered as the shoot, whereas the remaining portion comprising of the coco-peat plug embedded with the root hairs was considered as the “root-plug”. Fresh weight (FW) of the shoots and root-plugs was measured immediately, followed by oven drying at 85 °C for 48 h to measure the dry weight (DW). Subsequently, each root-plug was soaked in tap water for an hour, and the weight of water-saturated plugs was also recorded. Shoot water content and relative water content of root-plugs were calculated as follows:

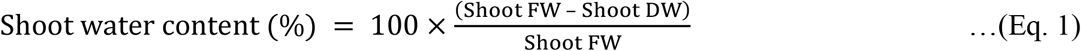

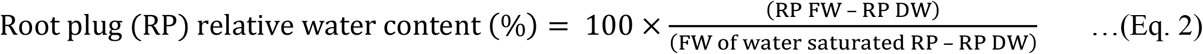

**Table 1.**
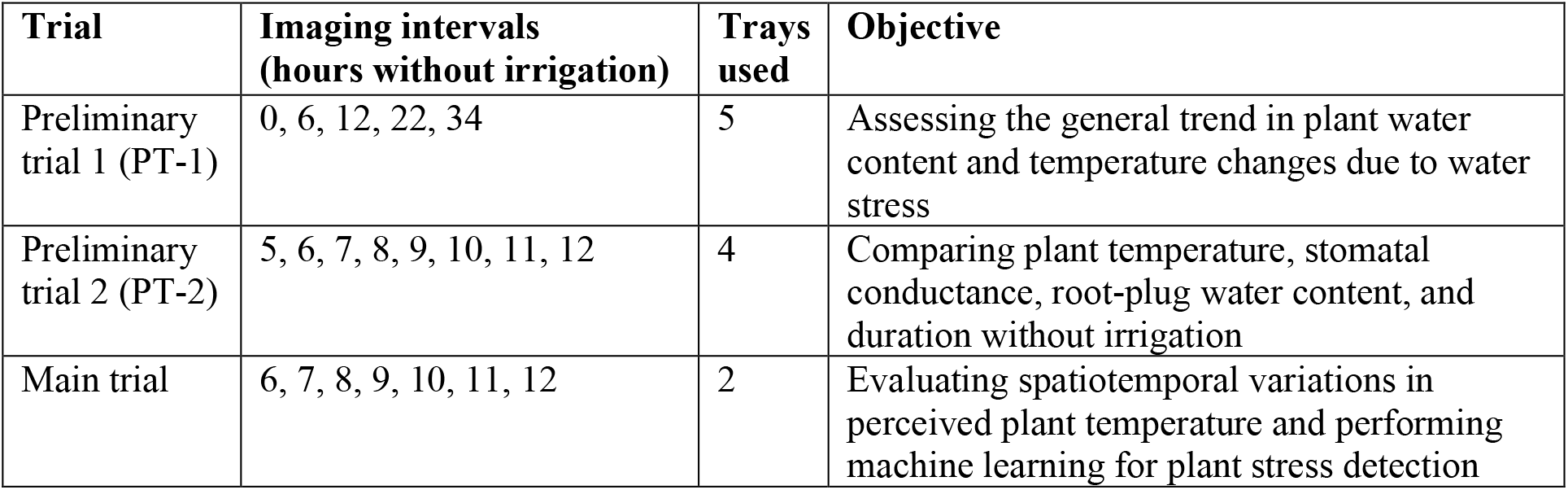
Overview of experimental design.

| **Trial** | **Imaging intervals**<br>**(hours without irrigation)** | **Trays**<br>**used** | **Objective** |
| --- | --- | --- | --- |
| Preliminary trial 1 (PT-1) | 0, 6, 12, 22, 34 | 5 | Assessing the general trend in plant water content and temperature changes due to water stress |
| Preliminary trial 2 (PT-2) | 5, 6, 7, 8, 9, 10, 11, 12 | 4 | Comparing plant temperature, stomatal conductance, root-plug water content, and duration without irrigation |
| Main trial | 6, 7, 8, 9, 10, 11, 12 | 2 | Evaluating spatiotemporal variations in perceived plant temperature and performing machine learning for plant stress detection |

The next trial (PT-2) was performed to better understand the responses at early stages of stress by comparing plant temperature with stomatal conductance and root-plug water content. As before, irrigation was stopped at 8 AM, and thermal imaging was performed using four trays at eight hourly intervals from 5 to 12 h without irrigation, wherein each tray was used for two random intervals (Table 1). Immediately after imaging, five plants were selected from the respective tray for recording stomatal conductance using an AP4 Porometer (Delta-T Devices, Cambridge, UK), and relative water content of root-plugs was calculated as before.

Findings of PT-1 and PT-2 were used to design the main trial. In this trial, thermal imaging of unirrigated plants was performed for two horizontally adjacent trays at hourly intervals from 6 to 12 h without irrigation (Table 1). This data was used for assessing spatiotemporal variations in perceived plant temperature, followed by ML for stress detection.

### 2.3 Thermal imaging

Thermal imaging was performed using a T1030sc thermal camera (Teledyne FLIR LLC, USA), with 7.5–14 *µ*m spectral range and focal plane array uncooled microbolometer with HD detector (resolution: 1024×768 pixels). The camera acquired thermal and RGB images simultaneously. Plants were imaged outside the vertical growing system using a customised setup for maintaining a fixed vertical distance (∼2 m) between the camera and the tray surface (Figure 1C), adjacent to the growth chamber to ensure similar environmental conditions. Imaging was performed under a neutral-white LED light source, at a room temperature of 25±1 °C. Camera parameters such as reflected, atmospheric, and optics temperatures were fixed (Prashar and Jones, 2016). Customised black body temperature reference pads were placed on each tray for the entire duration of the experiment (Prashar and Jones, 2016). Each tray was imaged within 30 sec of being taken out of the growth chamber, and was returned immediately.

### 2.4 Image pre-processing and temperature extraction

Pre-processing of thermal images was performed using Python programming (www.python.org) implementing the *flirextractor* module (https://pypi.org/project/flirextractor). The pipeline for this involved the following steps: 1) parallax correction between the thermal and RGB images for background (tray) removal using colour thresholding; 2) temperature mapping (15–30 °C scale) to grayscale (0–1 scale); 3) isolation of individual plants using an 8×8 grid to get 64 regions of interest (size: 95×95 pixels); 4) removal of unusable images; and 5) temperature extraction, followed by normalisation as an environmental correction for absolute errors (Figure 2A). Average plant temperature was computed after excluding a region of 10 pixels on all four sides of each plant to minimise the effect of overlapping leaves from adjacent samples (Figure 2B). The difference between observed plant temperature and reference temperature (RT), i.e., mean temperature of the black body reference pads, was used along with a constant (25 °C) to obtain normalised sample temperature as follows:

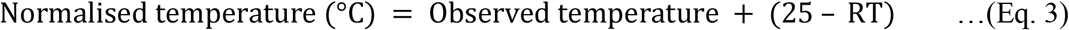

**Figure 2.**
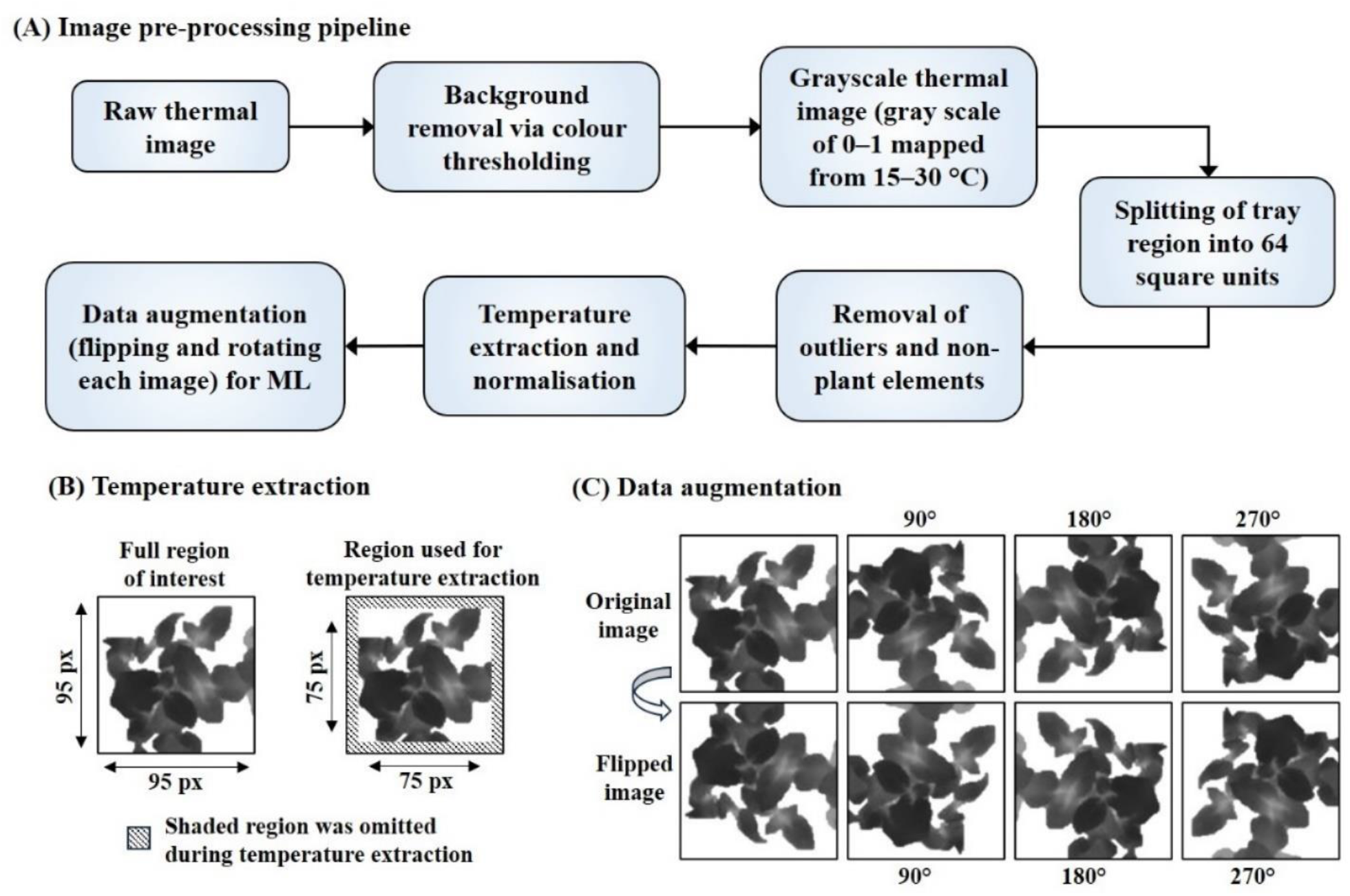
Thermal image pre-processing pipeline (A), data augmentation by flipping and rotating images (B), and selection of region of interest for extracting plant temperature. px, pixels.

Positional effects on perceived plant temperature within different regions of each tray were assessed by combining the data for both trays (Supplementary Figure S1) to get the average sample temperature at different positions.

### 2.5 ML for predicting plant stress

Thermal images from the main trial were used to design a Python-based ML pipeline for identifying stressed plants using the Scikit-learn module for SVMs (Pedregosa et al., 2011). Six ML models were generated for binary classification (BC), along with one model each for ternary and quaternary classification (TC, QC). In order to simplify the depiction of ML analyses, samples from each imaging interval have been represented using incremental stress levels (SL) as SL0, SL1, SL2, SL3, SL4, SL5, and SL6 for 6, 7, 8, 9, 10, 11, and 12 h without irrigation, respectively. Based on the results of PT-1 and PT-2, samples from 6 h without irrigation, i.e., SL0 class, were considered as “unstressed”. Each BC (two-class) model was trained using SL0 images along with images from one of the six other classes (SL1–SL6) as the stressed sample set. Models were labelled as BC-1 to BC-6, respectively. Similarly, the TC (three-class) and QC (four-class) models were generated using images from SL0, SL3, and SL6 (representing no, medium, and high stress, respectively) and images from SL0, SL2, SL4, and SL6 (representing no, mild, moderate, and high stress, respectively). The number of images used in each model has been summarised in Table 2.

**Table 2.**
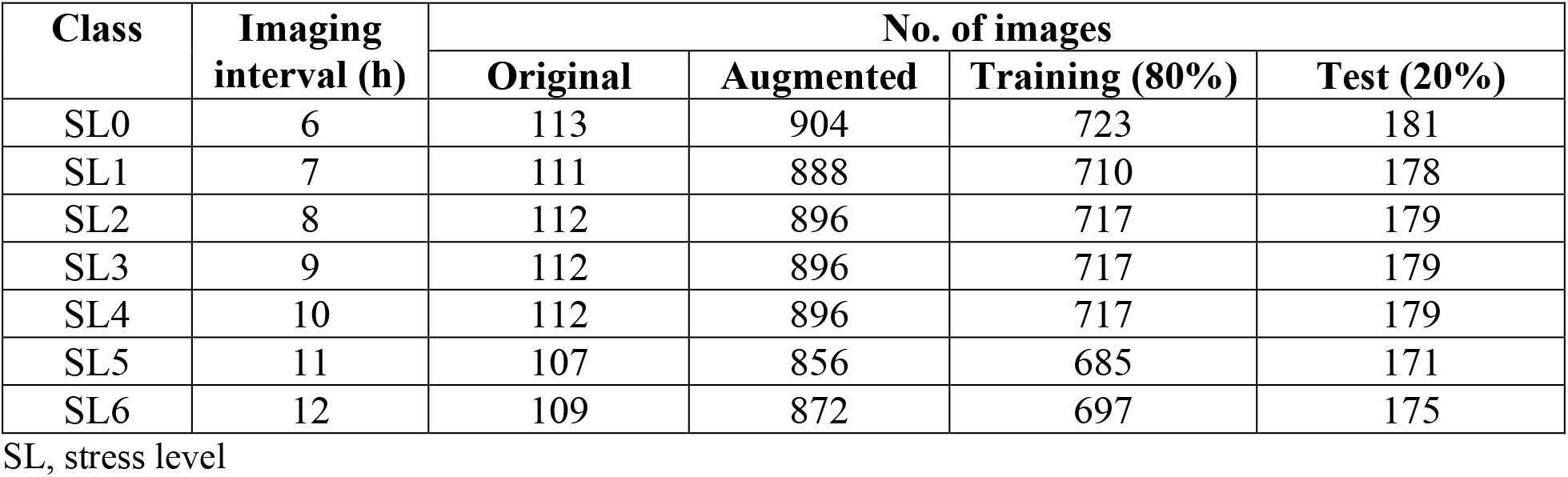
Summary of machine learning dataset.

Eight-fold data augmentation was performed by rotating and inverting each image (Figure 2C). This created a large training dataset for ML and helped bypass limitations associated with sample orientation. Subsequently, each thermal image was flattened to a vector containing the values for each pixel within the image. A randomised stratified train-test split of 80:20 was implemented. All eight ML models were allowed to choose from three kernels, viz., *linear* (*lin*), *polynomial* (*poly*), and *radial basis function* (*rbf*), along with a broad range of hyperparameters *C* (0.01–10) and *γ* (0.001–1) with the target of maximising accuracy. Here, *C* is the cost parameter defining penalty weight of deviations, and *γ* is an *rbf*-specific parameter controlling the trade-off between bias errors and variance in the adjusted model. The same hyperparameter sets were used for five-fold cross-validation to check model performance for variations in training datasets and hyperparameter combinations. Since the aim of the study was to develop a general understanding of the predictive capacity for different types of models from a plant science perspective, and not model optimisation, further hyperparameter tuning and validation was not done. Model performance was assessed by using the test dataset (20%) to calculate accuracy, *precision, recall* (sensitivity), *F1 score*, and *specificity* (Table 3) using the confusion matrix. The predictive performance of all ML models was further tested by individually deploying each model to analyse all samples from SL0–SL6 (Table 4), followed by plotting sample temperature against its Platt probabilistic estimate of stress provided by the respective ML model.

**Table 3.** Model performance parameters calculated from the confusion matrix.

| Parameter | Formula | Remarks |
| --- | --- | --- |
| Accuracy (%) | $100 \times \text{Total no. of TP for all groups} \div \text{Total no. of samples}$ | Indicates the overall performance of the model considering all classes |
| *Precision* | $\text{TP} \div (\text{TP} + \text{FP})$ | Share of correct classifications within each predicted class |
| *Recall* | $\text{TP} \div (\text{TP} + \text{FN})$ | Proportion of correctly classified samples from each original class |
| *F1 score* | $2 \times \text{Precision} \times \text{Recall} \div (\text{Precision} + \text{Recall})$ | Parameter indicating classification ability of the model for each class |
| *Specificity* | $\text{TN} \div (\text{TN} + \text{FP})$ | Goodness of model at identifying negative outcomes for each class |

FN, False negative; FP, False positive; TN, True negative; TP, True positive

**Table 4.** Summary of Platt’s probabilistic prediction dataset.

| Model | Known images |  | Unknown images | Total images |
| --- | --- | --- | --- | --- |
|  | Train | Test |  |  |
| BC-1 | 179 | 45 | 552 | 776 |
| BC-2 | 180 | 45 | 551 | 776 |
| BC-3 | 180 | 45 | 551 | 776 |
| BC-4 | 180 | 45 | 551 | 776 |
| BC-5 | 176 | 44 | 556 | 776 |
| BC-6 | 177 | 45 | 554 | 776 |
| TC | 267 | 67 | 442 | 776 |
| QC | 357 | 89 | 330 | 776 |

BC, binary classification model; TC, ternary classification model; QC, quaternary classification model

### 2.6 Statistical analysis

All data pertaining to shoot and root-plug water content as well as plant temperature were subjected to one-way ANOVA followed by Tukey’s post hoc test (*p* < 0.05) in Python using the *scipy*.*stats* library (https://docs.scipy.org/doc/scipy/reference/stats.html) to ascertain the significance of differences.

## Results

### 3.1 Plant stress response

Data from the first preliminary experimental trial (PT-1) revealed a gradual decline in shoot and root-plug water content along with a steady increase in plant temperature from 0 to 34 h without irrigation (Supplementary Figure S2). Slight decrease in mean shoot water content was observed from 6 to 12 h (Supplementary Figure S2A), whereas a significant reduction in root-plug water content and an increasing trend in plant temperature was perceptible over the same interval (Supplementary Figure S2B, C). Since a clear increasing trend in plant temperature along with a steep decline of *ca*. 74% reduction in root-plug water content was recorded between 0 to 12 h, the interval up to 12 h was targeted for IRT-based stress detection in the subsequent trials.

PT-2 data for 5 to 12 h without irrigation indicated that reduction in root-plug water content below ∼35% and stomatal conductance below ∼0.25 cm/s was accompanied with a steady rise in plant temperature (Figure 3A, B). Moreover, the threshold of 35% root-plug water content tentatively corroborated with 7 h without irrigation (Figure 3C). Root-plug water content loss slowed down from ∼15%/h between 5–7 h without irrigation to ∼2.6%/h between 7–12 h (Figure 3C), indicating reduced water uptake and stomatal closure starting from 7 h without irrigation, resulting in concomitant temperature rise and onset of stress. This suggests that plants were unstressed till 6 h without irrigation, followed by a gradual transition to stress from 6 to 7 h, and increasing levels of stress from 7 to 12 h.

**Figure 3.**
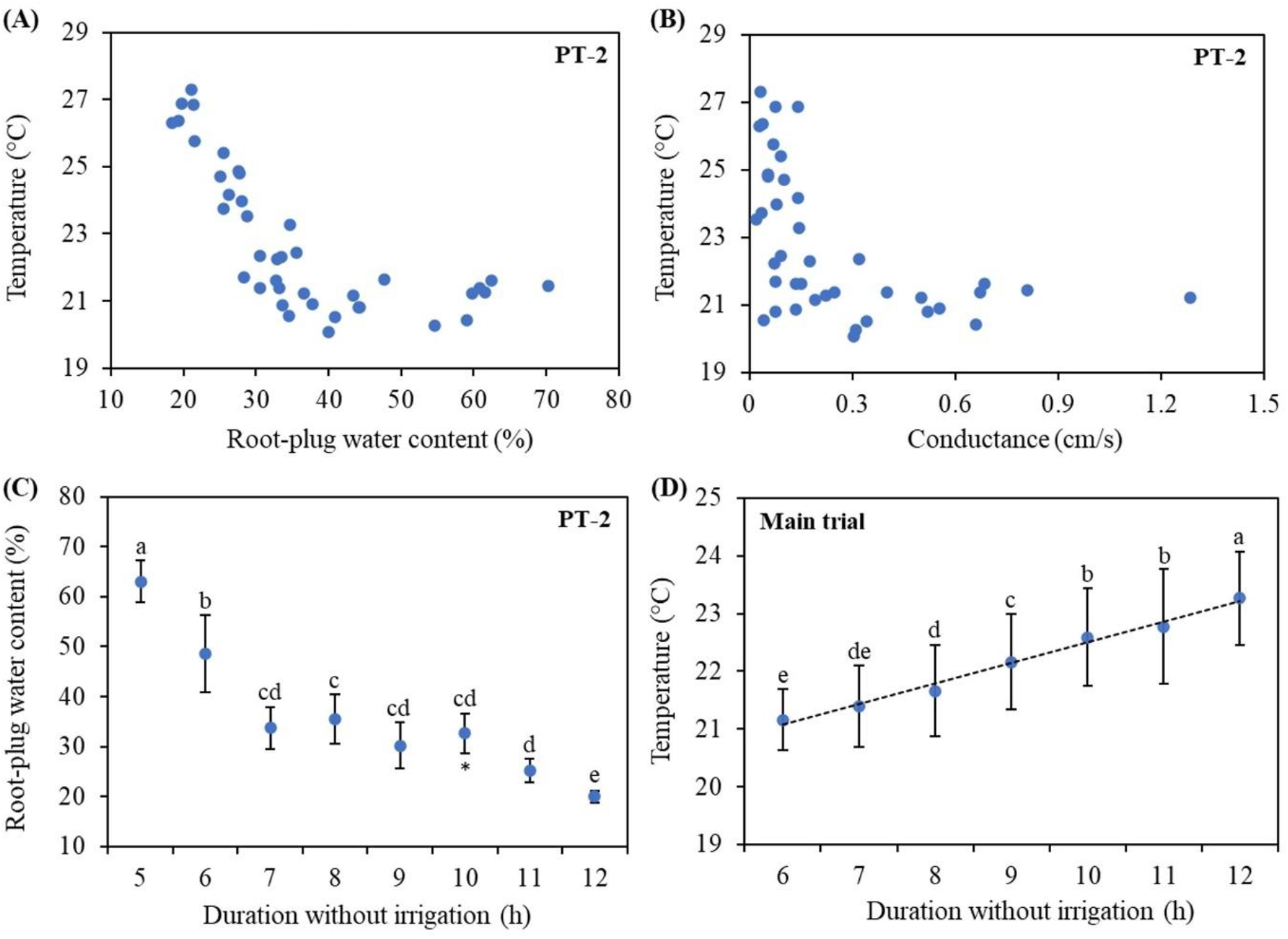
Relation of plant temperature with root-plug water content (A; *n* = 39) and stomatal conductance (B; *n* = 39), and root-plug water content at different intervals without irrigation (C; *n* = 5) recorded in preliminary trial 2 (PT-2), as well as plant temperature from 6 to 12 h without irrigation (D; *n* = 116) in the main trial. Error bars indicate mean ± standard deviation. Values indicated with different letters indicate significant difference in means as per Tukey’s HSD test (*p* < 0.05). *One outlier had to be omitted from PT-2 at 10 h due to handling error.

In the main trial, spatiotemporal trends in plant temperature were monitored prior to ML. A significant increase of *ca*. 2 °C (*p* < 0.05) in average plant temperature was noted from 6 to 12 h without irrigation (Figure 3D), corroborated by PT-1 data (Supplementary Figure S2C). Notably, a clear regional effect was perceptible within the tray area (Figure 4) wherein the samples closer to the air circulation vent always appeared cooler than the distant edge despite the level of stress being similar in terms of duration without irrigation. Further, higher temperatures were recorded only close to the center of the tray at later stages, indicating the likelihood of cooling effect due to better air circulation near all four edges.

**Figure 4.**
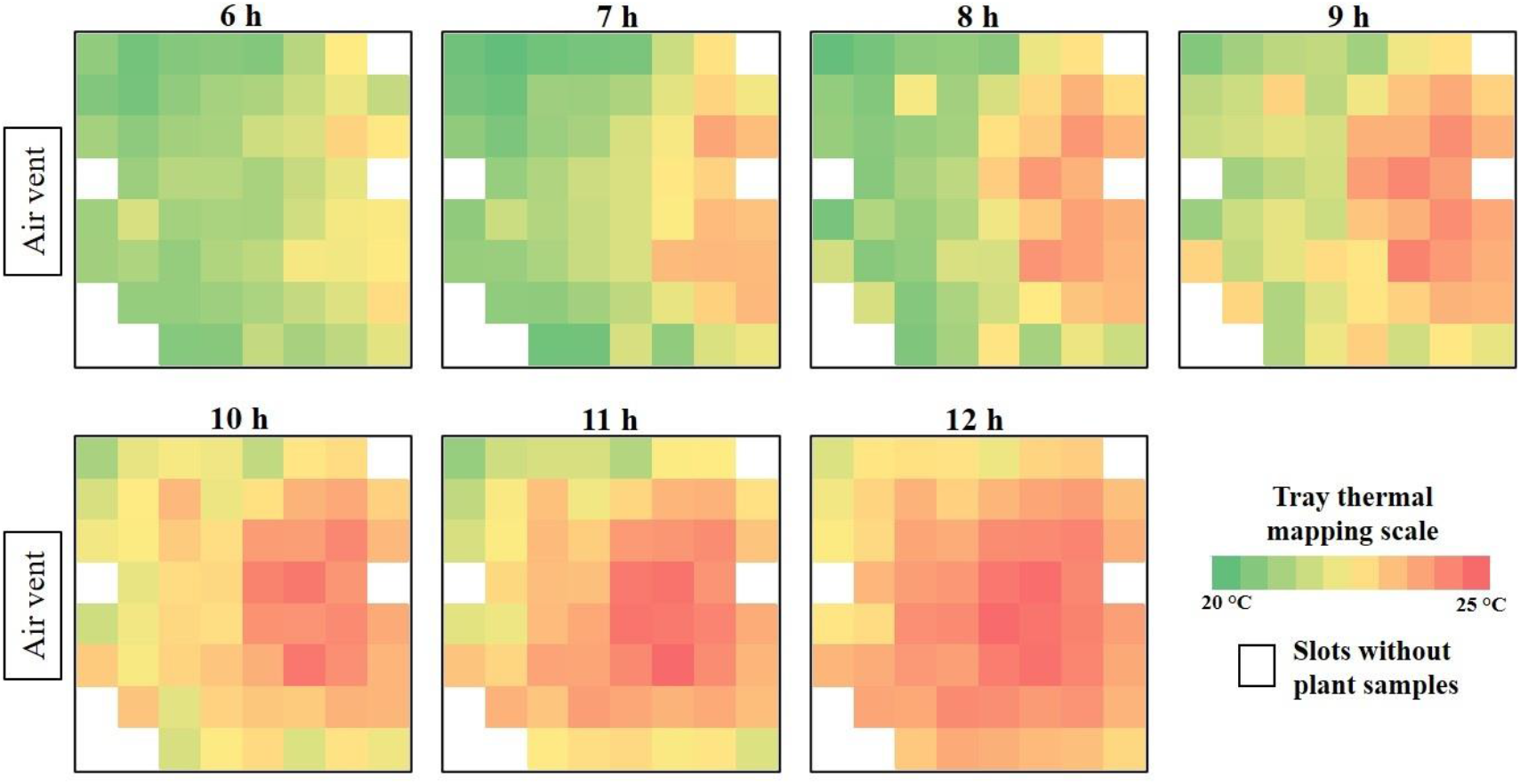
Heatmaps depicting spatial variation in temperature within the trays from 6 to 12 h without irrigation. Each image represents the average normalised temperature from two replicated trays from the main trial.

### 3.2 ML model performance

Training images were used for generating ML classification models (Table 2), whereas the confusion matrix for each model (Supplementary Tables S1, S2) was generated using the test dataset. Classification report for the BC models (Table 5) indicates an increase in accuracy and *precision* by *ca*. 1.7 times from BC-1 to BC-6 as the effect of stress intensified from 7 to 12 h. In contrast, although *recall, F1 score*, and *specificity* for SL0 (unstressed class) increased by *ca*. 1.25–1.5 times from BC-1 to BC-6, the same parameters showed an increment of *ca*. 2.3–2.9 times for the stressed classes. The TC model had higher accuracy compared to BC-1 and BC-2 models, whereas the QC model had the lowest overall accuracy (Tables 5, 6). *Precision, recall, F1 score*, and *specificity* of the TC model were higher for the two extreme classes, i.e., SL0 and SL6, as compared to the intermediate class, viz., SL3. A similar trend was observed for the QC model, wherein the magnitude of these four performance parameters was higher for SL0 and SL6 (extreme classes) as compared to SL2 and SL4 (intermediate classes). Five-fold cross-validation yielded similar accuracy values as the respective models (Supplementary Table S3), indicating consistency in model performance with variations in training and test datasets.

**Table 5.** Classification report on test data for binary classification (BC) models.

| Model | Class |  | Accuracy (%) | Precision |  | Recall |  | F1 score |  | Specificity |  | Support |  |
| --- | --- | --- | --- | --- | --- | --- | --- | --- | --- | --- | --- | --- | --- |
|  | UC | SC |  | UC | SC | UC | SC | UC | SC | UC | SC | UC | SC |
| BC-1 | SL0 | SL1 | 53.20 | 0.53 | 0.55 | 0.74 | 0.32 | 0.61 | 0.40 | 0.74 | 0.33 | 181 | 178 |
| BC-2 | SL0 | SL2 | 63.05 | 0.59 | 0.77 | 0.90 | 0.36 | 0.71 | 0.49 | 0.89 | 0.37 | 181 | 179 |
| BC-3 | SL0 | SL3 | 73.33 | 0.70 | 0.78 | 0.82 | 0.64 | 0.76 | 0.71 | 0.82 | 0.65 | 181 | 179 |
| BC-4 | SL0 | SL4 | 81.94 | 0.84 | 0.80 | 0.80 | 0.84 | 0.82 | 0.82 | 0.79 | 0.84 | 181 | 179 |
| BC-5 | SL0 | SL5 | 88.92 | 0.88 | 0.90 | 0.91 | 0.87 | 0.89 | 0.88 | 0.90 | 0.88 | 181 | 171 |
| BC-6 | SL0 | SL6 | 93.54 | 0.94 | 0.93 | 0.93 | 0.94 | 0.94 | 0.93 | 0.93 | 0.94 | 181 | 175 |

UC: Unstressed class; SC: Stressed class

**Table 6.** Classification report on test data for ternary and quaternary classification (TC, QC) models.

| Model | Class | Precision | Recall | F1 score | Specificity | Support |
| --- | --- | --- | --- | --- | --- | --- |
| TC | SL0 | 0.68 | 0.85 | 0.75 | 0.92 | 181 |
|  | SL3 | 0.54 | 0.32 | 0.40 | 0.66 | 179 |
|  | SL6 | 0.72 | 0.83 | 0.77 | 0.92 | 175 |
|  | *Model accuracy: 66.54%* |  |  |  |  |  |
| QC | SL0 | 0.49 | 0.82 | 0.62 | 0.94 | 181 |
|  | SL2 | 0.16 | 0.02 | 0.03 | 0.67 | 179 |
|  | SL4 | 0.39 | 0.41 | 0.40 | 0.80 | 179 |
|  | SL6 | 0.63 | 0.75 | 0.69 | 0.92 | 175 |
|  | *Model accuracy: 49.86%* |  |  |  |  |  |

### 3.3 Probabilistic prediction of plant stress via ML models

All ML models was individually deployed to analyse all 776 images from SL0 to SL6 (Table 4). Plots depicting plant temperature vs. the probability of sample being classified as “stressed” by the different models are shown (Figure 5, 6). In general, plants with higher temperatures had higher probability of being detected as stressed. Notably, models BC-3–BC-6 were able to distinguish between stressed and unstressed plants more robustly than BC-1 and BC-2, i.e., models trained using low-stress samples. A gradual transition in the distribution of the temperature-probability point cloud is visible across the models, from being localised linearly in the 0.4–0.65 probability region for BC-1 (Figure 5A) to that showing two discrete clusters in the 0 and 1 regions for samples with temperatures <22 °C and >23 °C, respectively, for the BC-6 model (Figure 5F). In concurrence with the BC models, the TC and QC models also assigned relatively cooler plants higher probabilities of being classified as SL0 or “unstressed” (Figure 6A, D). However, probabilistic predictions for higher levels of stress showed unclear trends in both TC and QC models (Figure 6B, C, E, F).

**Figure 5.**
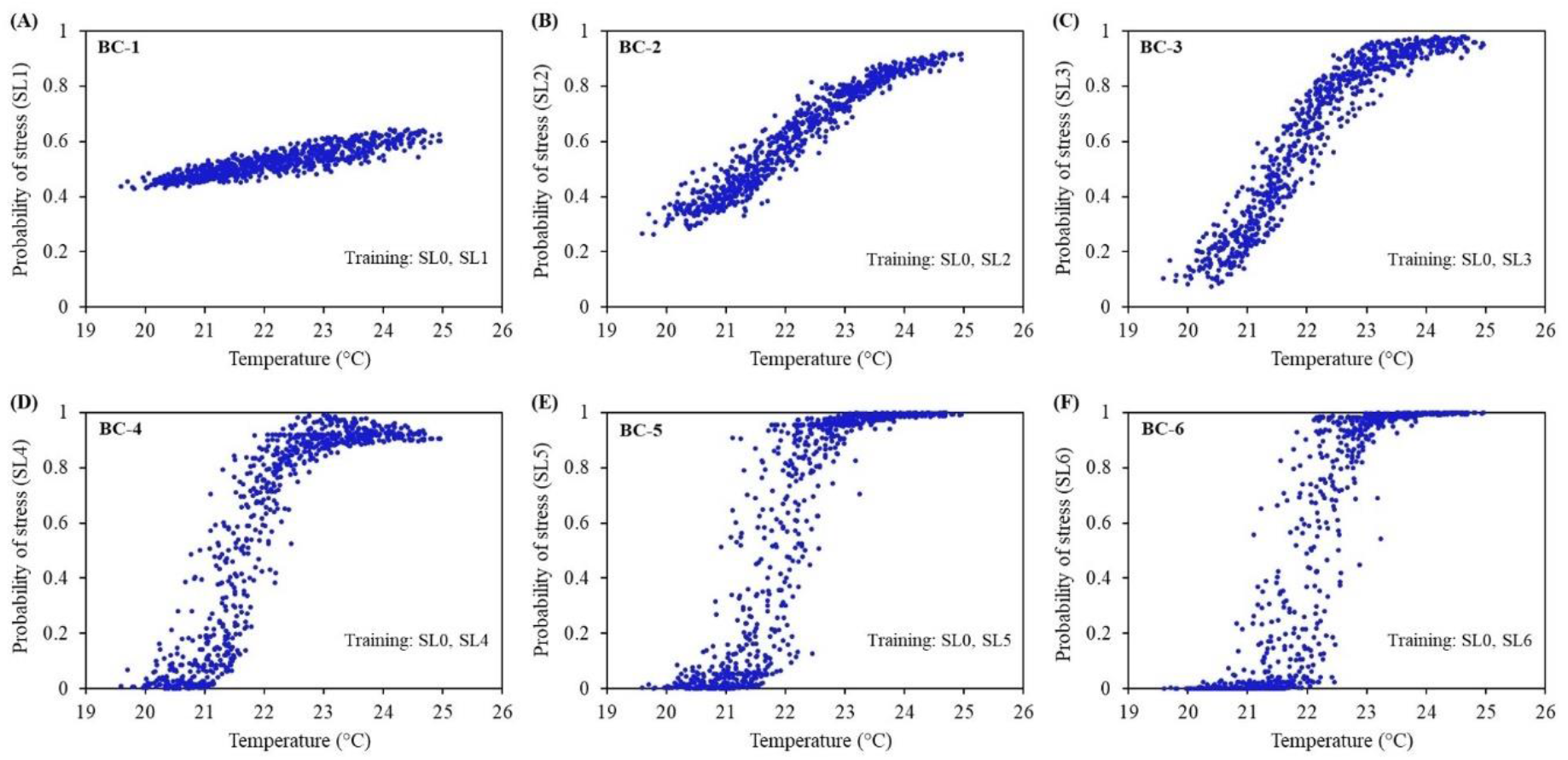
Plant temperature versus Platt’s probability of being classified as stressed by the six binary classification (BC) models (*n* = 776). Each plot represents the probabilistic prediction for all experimental samples. Stress level (SL) classes employed for creating the respective ML models have been specified within each plot along with the name of the model.

**Figure 6.**
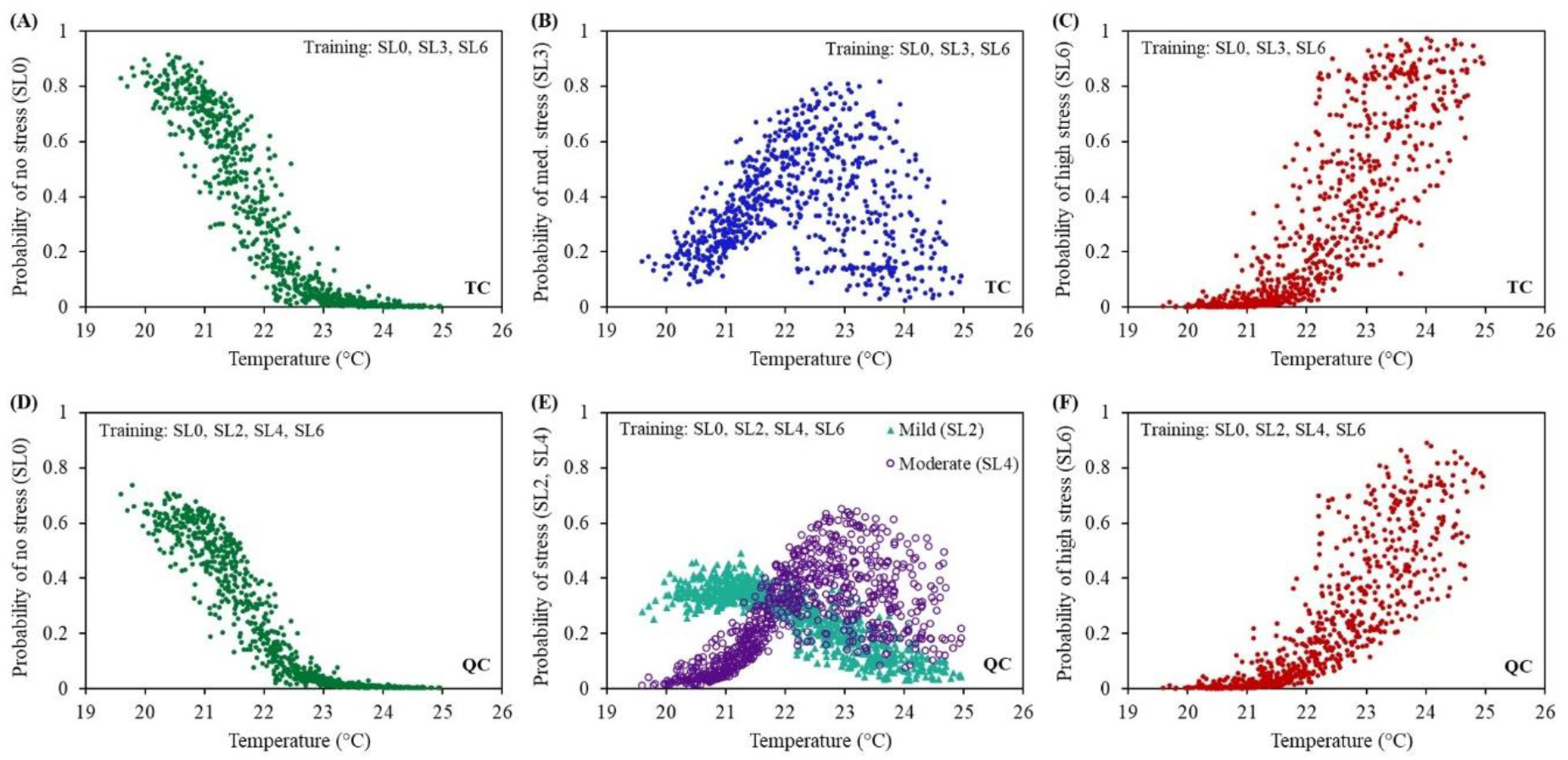
Plant temperature versus Platt’s probability of being classified into different levels of stress by the ternary and quaternary classification (TC, QC) models (*n* = 776). Each plot represents the probabilistic prediction for samples from all intervals. Stress level (SL) classes employed for creating the respective ML models have been specified within each plot along with the name of the model.

## 4 Discussion

In PT-1, although a significant reduction in relative water content of the root-plugs, i.e., from 95.5% to 49.8%, was observed from 0 to 6 h without irrigation, shoot water content and plant temperature remained relatively unchanged (*p* > 0.05) over this interval (Supplementary Figure S2). This suggests that the moisture content of root-plugs was high enough not to trigger temperature increase due to stomatal closure. However, subsequent reduction in plug water content from 49.8% to 22.8% over the 6–12 h interval (Supplementary Figure S2A) was accompanied with a decline in mean shoot water content by ∼12% (Supplementary Figure S2B) and a concomitant increase in average plant temperature by *ca*. 2 °C (Supplementary Figure S2C). This steady increase in plant temperature marked the triggering of stomatal closure resulting in reduced transpirational cooling (Jones, 1999; Grant et al., 2006; Ben-Gal et al., 2009), and the onset of stress.

In earlier studies, a significant increase in canopy temperature was reported for plants grown at *ca*. 50% field capacity compared to well-irrigated plants (Biju et al., 2018; Pradawet et al., 2023). In contrast, stability in canopy temperature observed in this study from 0 to 6 h without irrigation in PT-1 may be attributed to the stable RH and adequate moisture in the root-plug, possibly resulting in relatively stable vapor pressure deficit in the growth environment which delayed stomatal closure (Driesen et al., 2020). Co-assessment of root-plug water content, stomatal conductance, and plant temperature in PT-2 revealed that reduction in plug water content below a certain threshold, i.e., *ca*. 35%, triggered gradual stomatal closure and consequently caused a steady increase in plant temperature (Figure 3A, B). The connection between drought stress, stomatal closure, and increase in plant temperature is well-documented (Ben-Gal et al., 2009; Reynolds-Henne et al., 2010; Prashar and Jones, 2014), and forms the basis of the present inferences.

Based on the results of PT-1 and PT-2, the experimental window of 6 to 12 h without irrigation was deemed ideal for ML-based stress detection in the main trial. As seen here, an almost linear increase in average plant temperature was recorded from 6 to 12 h without irrigation, albeit with high variance at each interval (Figure 3D). This variability was mainly due to system design and the location of each plant. For example, plants located closer to the air circulation vents and edges were relatively cooler (Figure 4), possibly due greater air circulation and convective heat dissipation. Such observations mark the bottlenecks in identifying stressed plants within vertical farms solely based on absolute temperature thresholds, necessitating a more in-depth analysis of thermal images by ML to identify cohort-based probabilistic trends.

Performance parameters for the eight ML models indicated that accuracy of classification was higher when classes with more distinct levels of stress were used to train the models (Tables 5, 6). The trend was confirmed by model accuracy values obtained via five-fold cross-validation of each model (Supplementary Table S3). In general, ML models trained using adjacent or closer SLs, such as BC-1, BC-2, and QC models, had lower accuracy than the models whose training datasets were further apart (Tables 5, 6). This may be attributed to similarities in stress response, i.e., change in canopy temperature, between classes with similar levels of stress. In addition, positional effects within the tray (Figure 4) increased temperature variance for each SL, further reducing the discernability of samples between closer SLs.

Each of the five performance parameters (Table 2) derived from the confusion matrix represented a unique aspect of model “confusion”, i.e., the ability or inability to distinguish two classes robustly. While “accuracy” provided a general overview of model performance, the other four parameters provided class-specific insights on model behaviour. For instance, BC-1 had similar *precision* for both SL0 (0.53) and SL1 (0.55), although the *recall* was 0.74 and 0.32 for the two classes, respectively (Table 5). This indicates that almost half of the samples categorised as SL0 or SL1 were classified correctly by the BC-1 model, whereas more samples originally belonging to SL0 were correctly identified than SL1. The *F1 score* generalised this inference by combining the outcomes of the previous two parameters. In consensus, the low *specificity* score for SL1 (Table 5) suggests that the BC-1 model was also unable to reliably identify samples that did not belong to SL1, whereas it was able to correctly recognise most of the samples (74%) that did not belong to SL0. Since SL0 and SL1 had somewhat similar levels of stress, it is very likely that the model was confused. Results for BC-2 were similar, albeit slightly better (Table 5).

Among the BC models, BC-3 showed the earliest possibility of reliable stress prediction by correctly identifying a majority (115/179; Supplementary Table S1) of the samples from the stressed class (SL3) with an overall accuracy of 73.33% (Table 5). The *recall* values for SL0 and SL3 were 0.82 and 0.64, respectively, indicating more samples from SL0 being identified accurately. *F1 score* for BC-3 indicates that the model could classify SL0 samples more reliably than SL3 samples (Table 5). This was further validated by the considerably higher *specificity* for SL0 (0.82) as compared to SL3 (0.65). Nonetheless, probability estimates for the likelihood of stress with respect to plant temperature for this model yielded a uniform and continuous distribution of points throughout the temperature range (Figure 5C), unlike the plots for BC-1 and BC-2 (Figure 5A, B) which showed limited probabilistic ranges. Furthermore, although the model was trained with only SL0 (unstressed) and SL3 (medium stress) samples, it was able to accurately predict a higher probability of stress (>0.8) for samples at >23 °C, mostly belonging to SL5 and SL6 (high stress).

The BC-4 model showed some similarities as BC-3 in predicting the probability of stress (Figure 5D). However, it exhibited an erratic trend at intermediate temperatures (∼21.5–22.5 °C), which was more prominent in BC-5 and BC-6 (Figure 5E, F). Specifically, probability distributions for plants at very low and high temperatures were respectively localised at the two extremities, i.e., close to 0 and 1, while the datapoints for intermediate temperatures were distributed arbitrarily between probability values of 0.1 and 0.9 for both models. Classification report for BC-4, BC-5, and BC-6 indicated >80% accuracy, whereas values for all other performance parameters ranged between 0.79–0.94 (Table 5). This suggests that these three models were able to reliably classify test images, which were similar to the “known” training samples, although “unknown” samples from intermediate stress levels, i.e., SL2 and SL3, might have confused the models during probabilistic prediction.

Results for the TC model revealed that it could distinguish SL0 samples from SL3 and SL6 reliably, but failed to distinguish between SL3 and SL6 effectively (Figure 6A–C). While a continuous distribution for probabilistic prediction of SL0 was observed with this model (Figure 6A), the prediction was relatively irregular when attempting to distinguish between medium (SL3) and high (SL6) levels of stress (Figure 6B, C). This could have been due to the higher variability and overlap in canopy temperature between later stages of stress. Moreover, despite an overall accuracy of 66.54%, all four class-specific performance parameters for TC (Table 6) indicated that the model was fairly confused while identifying SL3 samples, which were more reliably identified by BC-3 (Table 5). Since the TC model could be technically considered as an extension of the BC-3 model with an extra “high stress” class, viz., SL6, it may be inferred that having too many classes confused the model.

In the QC model, probabilistic classification values were below 0.7 for SL0, SL2, and SL4 (Figure 6D, E). This indicates that the model could not identify samples confidently for these classes. Moreover, probability distribution for SL4 and SL6 was highly inconsistent for this model (Figure 6E, F). This, along with an overall accuracy of <50%, may again be attributed to significant similarity between samples of adjacent classes and increased model confusion, similar to that observed in the TC model. Thus, the outcomes of TC and QC models highlight the limitation of SVM-based thermal image classification models trained with too many classes and samples having relatively high degree of similarity.

Since the BC-3 model showed a steady probabilistic transition over the entire temperature range and was even able to reliably assess the likelihood of stress in samples beyond its training range (Figure 5C), such a model could be deemed useful for predicting the degree of stress over a continuous scale in a probabilistic manner rather than binary classification as “stressed” or “unstressed”. This could be especially beneficial in real-world scenarios where the level of plant stress needs to be assessed with precision. Probabilistic prediction outcomes generated by using each model to analyse “known” and “unknown” images from all stress levels (Table 4) provided an insight on how each type of model would perform in a real-world situation where the target plant samples could be at any level of stress. Furthermore, spatial variability in temperature at each level of stress due to positional effects (Figure 4) was helpful in designing more realistic and robust ML training models, as low variance datasets would increase the chances of overfitting during model training.

Earlier studies have explored the implementation of ML and IRT along with other imaging techniques for assessing crop stress in a variety of ways. For instance, Raza et al. (2014) used IRT in conjunction with ML employing thermal and colorimetric features, and implemented them in different combinations for probabilistic detection of water-deficit samples. Banerjee et al. (2018) explored the implementation of thermal imaging coupled with ML for estimating leaf area index in wheat exposed to drought, whereas Das et al. (2021) employed unmanned aerial vehicle-based IRT, and analysed wheat images using classification and regression trees to predict yield, with the final aim of identifying drought tolerant cultivars. In contrast, Correia et al. (2022) utilised multispectral images to create masks for segmenting thermal images of droughted wheat, followed by regression modelling for predicting plant biomass using image features.

The present study followed a distinct IRT pipeline for identifying stressed plants by combining the following elements: 1) use of colorimetric features for image segmentation, 2) direct deployment of thermal images for ML, 3) temporal thermal imaging for continuous monitoring of stress responses, and 4) probabilistic prediction of stress. The SVM algorithm used in this study not only considered the magnitude of values at each image pixel, but also accounted for the spatial distribution of data points for each sample. As the current dataset comprised of canopy images of individual plant samples, each image was used eight times by flipping and rotating (Figure 2B) to create eight unique versions of each image by rearranging the pixel values within the flattened vector. This data augmentation process helped overcome any potential biases arising from the spatial distribution of image pixels that were not perceptible visually, and created larger and more diverse training datasets, which is beneficial for reducing overfitting of ML models (Leng et al., 2017).

The variety of applications for IRT reported previously (Raza et al., 2014; Banerjee et al. 2018; Das et al., 2021; Correia et al., 2022) highlight the versatility of the technology for high-throughput real-time assessment of water stress and crop performance in field studies. The present study attempts to address a distinct topic, i.e., stress detection by IRT within vertical farms and its challenges, which has not received a lot of attention as yet. In general, the technique can be effectively translated for detecting any stress where plant temperature is affected, and where visual symptoms are not prominent at early stages, such as stress due to inadequate root aeration caused by overwatering or waterlogging, as well as biotic stresses where stomatal response is affected, such as root infections. Since research related to improving resource-use efficiency in vertical farms, especially light and water use efficiency, is being pursued actively, IRT-based crop monitoring could be helpful in such research areas by aiding researchers and growers to monitor plant status more accurately while testing new growth protocols.

Rapid growth in implementation of vertical farming systems over the past decade has revealed various relevant challenges, hence, revealing the scopes for improvement. Since such systems rely extensively upon technology with numerous mechanical components running continuously, any undetected incident resulting in plant stress may have a cascading effect causing extensive crop loss. Hence, early detection of plant stress plays a pivotal role in ensuring productivity in such highly-controlled and machine-dependent systems, prompting the adoption of IRT for such systems. The present results indicate that while longer durations of stress made it easier to identify stressed plants, even mild symptoms appearing early could be detected via IRT by using ML models. The present report also highlights the pitfalls of using absolute temperature for plant stress detection in vertical farms, emphasising the importance of spatial thermal mapping in such systems. Further investigations using other crops and different ML approaches with different stresses will broaden the knowledge base for implementing this technology in commercial vertical farms feasibly.

## Author contributions

NB, VACG, TH and AP conceived the project. AA, FdJC and AP designed the study. AA and FdJC performed the experiments with help from RD. AA performed image and data analysis with involvement from FdJC and RD on numerical data analysis. AA and AP drafted the manuscript and all authors contributed to amendments and revisions. All authors approved the submitted version.

## Acknowledgments

We thank the InFarm UK team for supplying seedlings and providing technical support, along with InFarm Crop Science team (Germany) for their support. We acknowledge all the partners (RoboScientific, Marks and Spencer, and InFarm) for their feedback and support in the project. We also thank the staff at Newcastle University for their technical, administrative and logistic support. AA thanks Dr. Monica Barman, Leibniz Institute for Vegetable and Ornamental Crops (IGZ), for her guidance on statistical analysis.

## Data availability statement

The datasets used in this study are available from the corresponding author upon reasonable request.

