## Supplementary material for "Infrared thermography for plant stress detection in vertical farms: Investigating spatiotemporal variations and exploring solutions via machine learning"

**Supplementary Table S1** Confusion matrices for the binary classification (BC) models.

| | | | | | | | |
|---|---|---|---|---|---|---|---|
| **BC-1** | | *Predicted* | | **BC-2** | | *Predicted* | |
| | | *SL0* | *SL1* | | | *SL0* | *SL2* |
| *Actual* | *SL0* | 134 | 47 | *Actual* | *SL0* | 162 | 19 |
| | *SL1* | 121 | 57 | | *SL2* | 114 | 65 |
| **BC-3** | | *Predicted* | | **BC-4** | | *Predicted* | |
| | | *SL0* | *SL3* | | | *SL0* | *SL4* |
| *Actual* | *SL0* | 149 | 32 | *Actual* | *SL0* | 144 | 37 |
| | *SL3* | 64 | 115 | | *SL4* | 28 | 151 |
| **BC-5** | | *Predicted* | | **BC-6** | | *Predicted* | |
| | | *SL0* | *SL5* | | | *SL0* | *SL6* |
| *Actual* | *SL0* | 164 | 17 | *Actual* | *SL0* | 169 | 12 |
| | *SL5* | 22 | 149 | | *SL6* | 11 | 164 |

SL, stress level used as the class for machine learning.

**Supplementary Table S2** Confusion matrices for the ternary and quaternary classification (TC, QC) models.

| | | | | | | | | | | |
|---|---|---|---|---|---|---|---|---|---|---|
| **TC** | | *Predicted* | | | **QC** | | *Predicted* | | | |
| | | *SL0* | *SL3* | *SL6* | | | *SL0* | *SL2* | *SL4* | *SL6* |
| *Actual* | *SL0* | 153 | 25 | 3 | *Actual* | *SL0* | 149 | 5 | 27 | 0 |
| | *SL3* | 67 | 58 | 54 | | *SL2* | 108 | 3 | 48 | 20 |
| | *SL6* | 5 | 25 | 145 | | *SL4* | 41 | 9 | 73 | 56 |
| | | | | | | *SL6* | 5 | 2 | 37 | 131 |

SL, stress level used as the class for machine learning.

**Supplementary Table S3** Model accuracy values obtained from five-fold cross-validation.

| **Model** | | **Accuracy (%)** | | | | | |
|---|---|---|---|---|---|---|---|
| **Name** | **Classes** | **Iteration 1** | **Iteration 2** | **Iteration 3** | **Iteration 4** | **Iteration 5** | **Mean ± SD** |
| BC-1 | SL0, SL1 | 57.38 | 57.66 | 57.54 | 56.98 | 59.22 | 57.76 ± 0.86 |
| BC-2 | SL0, SL2 | 61.11 | 60.00 | 63.61 | 64.72 | 65.00 | 62.89 ± 2.23 |
| BC-3 | SL0, SL3 | 72.50 | 73.33 | 74.72 | 73.89 | 75.28 | 73.94 ± 1.10 |
| BC-4 | SL0, SL4 | 81.11 | 83.06 | 84.44 | 82.22 | 81.94 | 82.55 ± 1.26 |
| BC-5 | SL0, SL5 | 85.80 | 89.20 | 90.63 | 87.78 | 89.77 | 88.64 ± 1.89 |
| BC-6 | SL0, SL6 | 90.17 | 92.39 | 95.49 | 95.21 | 95.49 | 93.75 ± 2.39 |
| TC | SL0, SL3, SL6 | 60.37 | 62.24 | 70.04 | 70.97 | 72.47 | 67.22 ± 5.51 |
| QC | SL0, SL2, SL4, SL6 | 47.76 | 51.68 | 53.36 | 50.35 | 52.59 | 51.15 ± 2.20 |

BC, Binary classification; TC, Ternary classification; QC, Quaternary classification; SL, stress level used as the class for machine learning.

### Supplementary figures

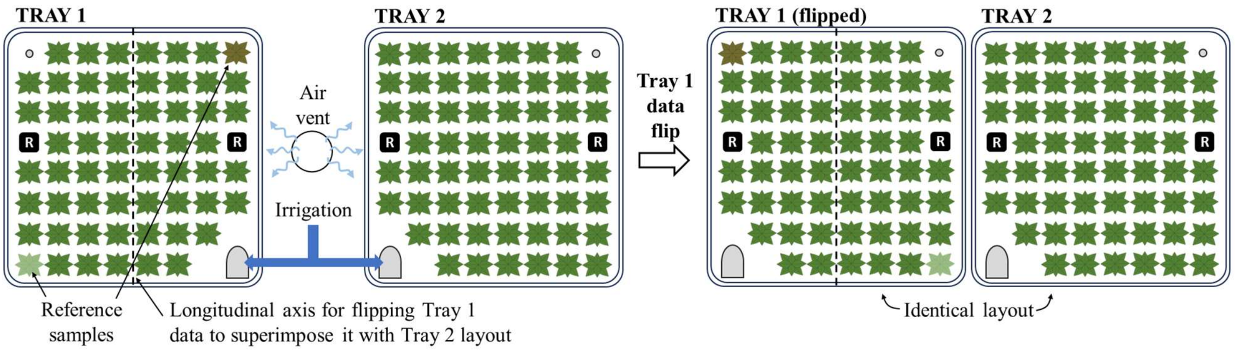

**Supplementary Fig. S1** Flipping of tray data for spatial superimposition. Trays 1 and 2 are replicate trays used in the main trial for machine learning with thermal images. Data for Tray 1 was flipped along the longitudinal axis to enable spatial superimposition with Tray 2 data for calculating the average temperature at each position within the tray relative to the edges, air vents, and irrigation supply. Two samples on Tray 1 have been shaded differently (reference samples) to understand the effect of flipping.

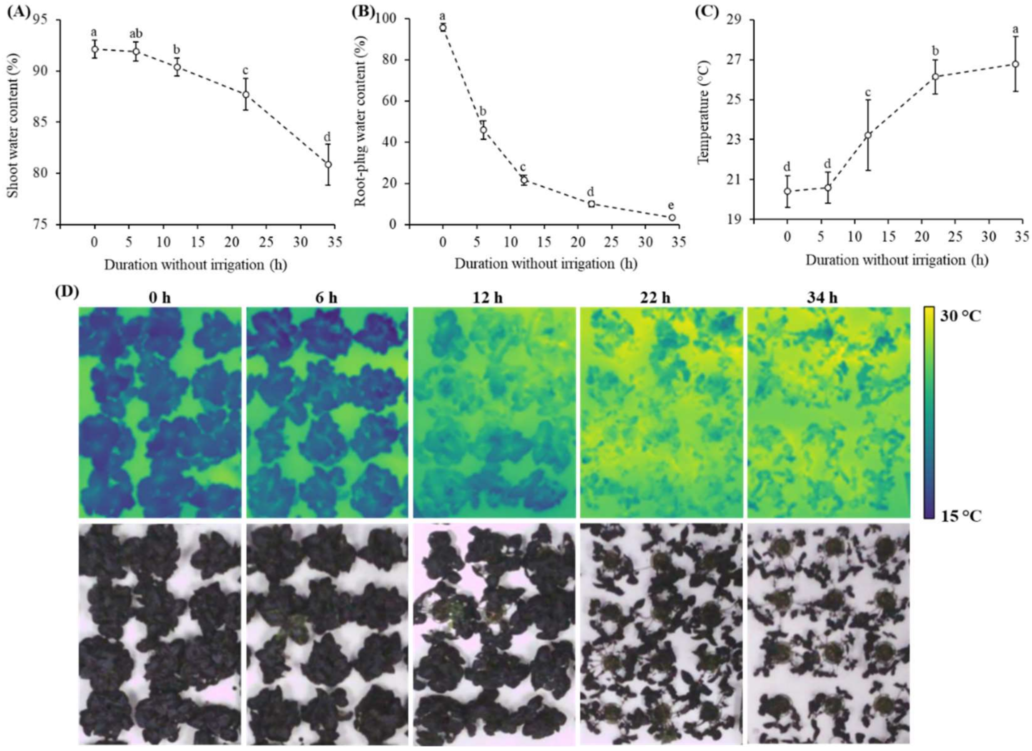

**Supplementary Fig. S2** Shoot (A) and root-plug (B) water content ($n$ = 12), plant temperature (C, $n$ = 94), and sample images (D; top: thermal images; bottom: colour images) from 0 to 34 h without irrigation as recorded in preliminary trial 1 (PT-1). Error bars indicate mean ± standard deviation. Values indicated with different letters indicate significant difference in means as per Tukey's HSD test ($p < 0.05$).
